# Forecasting cell state futures from static snapshots

**DOI:** 10.64898/2026.02.08.704720

**Authors:** Erpai Luo, Haoxiang Gao, Haiyang Bian, Shuguang Peng, Yaru Li, Chen Li, Minsheng Hao, Mo Chen, Yuli She, Lei Wei, Kai Liu, Xuegong Zhang

## Abstract

Predicting future cell state transitions from static transcriptomic snapshots is vital for understanding and steering cellular dynamics. Although existing trajectory and RNA velocity methods can infer local transitions within observed data, they are fundamentally limited in extrapolating long-range cellular dynamics. Here, we introduce CellTempo, an autoregressive generative framework that enables long-range forecasting of cell state trajectories directly from static transcriptomic snapshots. CellTempo is pretrained on scBaseTraj, a large-scale collection of multi-step cellular transition sequences constructed by integrating diverse experimental technologies and data sources. Across diverse biological systems, CellTempo accurately predicts long-range lineage progression and reconstructs unseen cellular potential landscapes. It further distinguishes transient perturbation responses from long-term cell-fate changes, enabling prioritization of candidate compounds for cell-fate engineering. These results demonstrate that long-range cellular dynamics are inferable from static observations, establishing a foundation for predictive and controllable modeling of cell fate.

## Introduction

Single-cell sequencing has enabled comprehensive mapping of cell state spaces across tissues, developmental processes, and disease contexts^1-8^. However, these maps are inherently static, because each measurement captures only a molecular snapshot at a single time point. As a result, they provide limited insight into how cellular states transition and what trajectories cells may adopt in the future.

A central question therefore arises: can static transcriptomic snapshots be used to forecast future state transitions of cells? Existing computational approaches have made substantial progress in extracting temporal structure from snapshot data. Methods based on pseudotime ordering^9^, RNA velocity^10^, and trajectory inference^11^ can interpolate transition events within the range of observed cells and infer immediate transition tendencies^12,13^. However, these approaches operate largely within the manifold defined by observed data, and whether static transcriptomes contain sufficient information to support long-range predictive modeling of cell state evolution remains unclear. This gap limits both our understanding of development and disease progression as well as our ability to design interventions that redirect cell fate.

Here we formulate cell-state evolution as a generative forecasting problem and present CellTempo, an autoregressive generative AI model designed to forecast long-range future cellular trajectories directly from static single-cell transcriptomes. CellTempo is pretrained on scBaseTraj, the first large-scale database of multi-step cellular transition sequences constructed by integrating multiple existing technologies and dynamic information. Through this pretraining, CellTempo learns latent dynamical regularities governing cell state progression, enabling forward-time forecasting without requiring external dynamic information at inference. Starting from a single observed cell, the model generates ordered sequences of future states, providing a probabilistic view of how cellular trajectories may unfold. This generative formulation further allows reconstruction of cellular potential landscapes that capture the preferred directions and tendencies of cell state progression. Within the same dynamical framework, CellTempo supports perturbation-conditioned forecasting, allowing it to distinguish transient transcriptional changes from sustained alterations in cell fate.

We demonstrate these capabilities across diverse biological systems. The long-range cellular trajectories generated by CellTempo faithfully recapitulate transcriptional dynamics, lineage topology, and cell-type composition observed in real systems. In human bone-marrow hematopoiesis, CellTempo reconstructs a cellular potential landscape that reliably reflects the structured progression of stem and progenitor populations. Within this landscape, targeted perturbation of lineage-specific gene modules systematically redirects hematopoietic stem cells toward distinct differentiation fates, including erythroid and lymphoid lineages. Applied to the cell-fate engineering of oligodendrocyte precursor cells (OPCs), CellTempo moved beyond perturbation models that primarily predict immediate transcriptional responses, prioritizing interventions by whether those responses ultimately developed into sustained OPC formation. These results demonstrate that static single-cell measurements encode predictive information about cellular dynamics beyond the observed state space, enabling long-range forecasting of whether immediate perturbation responses translate into sustained changes in cell fate.

Together, this work establishes a predictive approach to single-cell biology in which static transcriptomes are used not only to reconstruct observed trajectories but also to forecast future cell-state transitions, opening new opportunities to study development, disease progression, and therapeutic responses, as well as to rationally design interventions that reshape cell fate.

## Results

### CellTempo: an autoregressive AI model for future cellular trajectories forecasting

To model cellular dynamics over time, we propose CellTempo, a temporal single-cell AI model designed for autoregressive forecasting of long-range cell state evolution trajectories. Rather than treating cells as static observations to be ordered or embedded^14-16^, CellTempo formulates cellular dynamics as a long-term cellular trajectory. It employs an autoregressive strategy to learn the contextual relationships among cells within the trajectory, enabling future cell states to be generated conditioned on present observations (Figure 1A). CellTempo can directly predict the cellular trajectories from a single observed cell and provides a unified generative basis for trajectory forecasting, cellular potential landscape reconstruction, and perturbation response prediction (Figure 1B).

**Figure 1.**
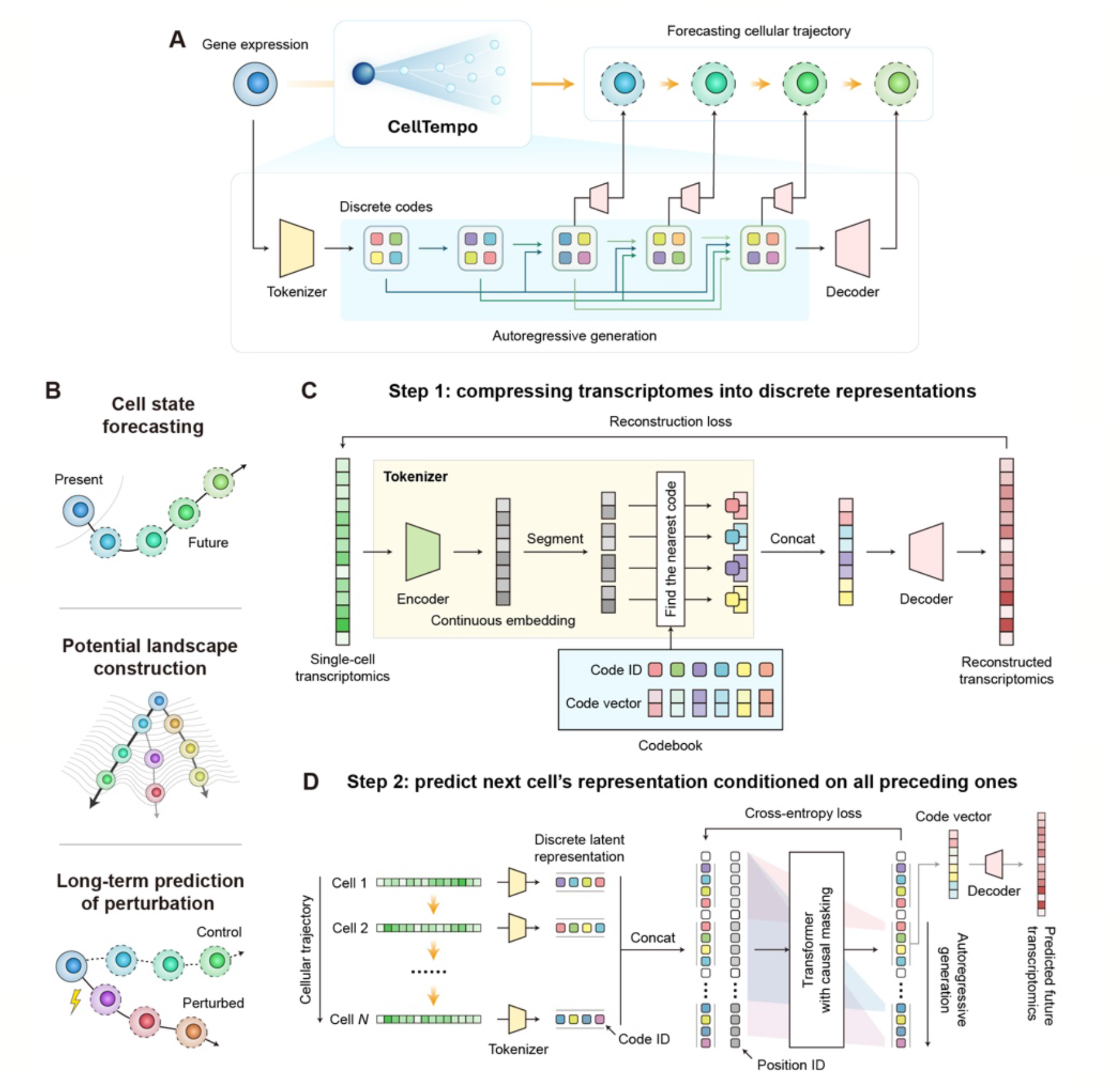
Overview of CellTempo. (A) Wrokflow of CellTempo. It tokenizes the transcriptome of a known cell into discrete latent representations, uses this known cell to predict the latent representations of future cell states, and decodes them back into gene expression. (B) Representative applications of CellTempo. (C-D) Training procedure and model architecture of CellTempo. In the first step (C), we train a cell VQ-VAE model to compress single-cell transcriptomes into discrete latent representation and reconstruct transcriptomes from these latents. In the second step (D), given cell trajectory data, we use the trained cell VQ-VAE tokenizer to obtain Code IDs of each cell’s latent representation and concatenated these Code IDs along the trajectory. These concatenated Code IDs, together with position IDs indicating cell indices in the trajectory, are fed into Transformer with causal masking to autoregressively predict the next cell’s latent representation.

We designed CellTempo with a two-stage architecture that decouples cellular representation learning from temporal sequence modeling. In the first stage, each single-cell transcriptome is compressed into a compact discrete representation using a vector-quantized variational autoencoder (VQ-VAE), substantially reducing dimensionality while preserving biologically meaningful information (Figure 1C, Methods). By transforming high-dimensional and noisy gene expression profiles into discrete latent codes, this representation provides a stable and compact substrate for autoregressive modeling of long-range cellular trajectories, while allowing accurate reconstruction back to the original expression space (Figure S1).

In the second stage, CellTempo models cellular dynamics using a transformer-based autoregressive generative model that operates on sequences of discrete cell representations (Figure 1D, Methods). Cells along a trajectory are encoded as temporally ordered sequences of discrete representations, forming trajectory-level sequences analogous to token sequences in natural language modeling. This model is trained with causal masking to predict the discrete representation of the next cell conditioned on all preceding ones, enabling the learning of long-range temporal dependencies across cell states. During inference, CellTempo starts from a single observed cell and autoregressively generates future states in latent space, which are subsequently decoded to obtain predicted gene expression profiles.

Together, this formulation and architecture establish CellTempo as a scalable model for autoregressive forecasting of cellular dynamics, forming the foundation for all subsequent analyses.

### Constructing large-scale trajectory datasets for training autoregressive forecasting models

Training autoregressive models of cellular dynamics requires multi-step cellular transition sequences, but such supervision is rarely available in single-cell studies. We therefore constructed the first large-scale single-cell trajectory dataset, scBaseTraj, which integrates RNA velocity, pseudotime, and trajectory inference methods to transform static snapshots into biologically grounded multi-step cellular sequences (Figure 2A).

**Figure 2.**
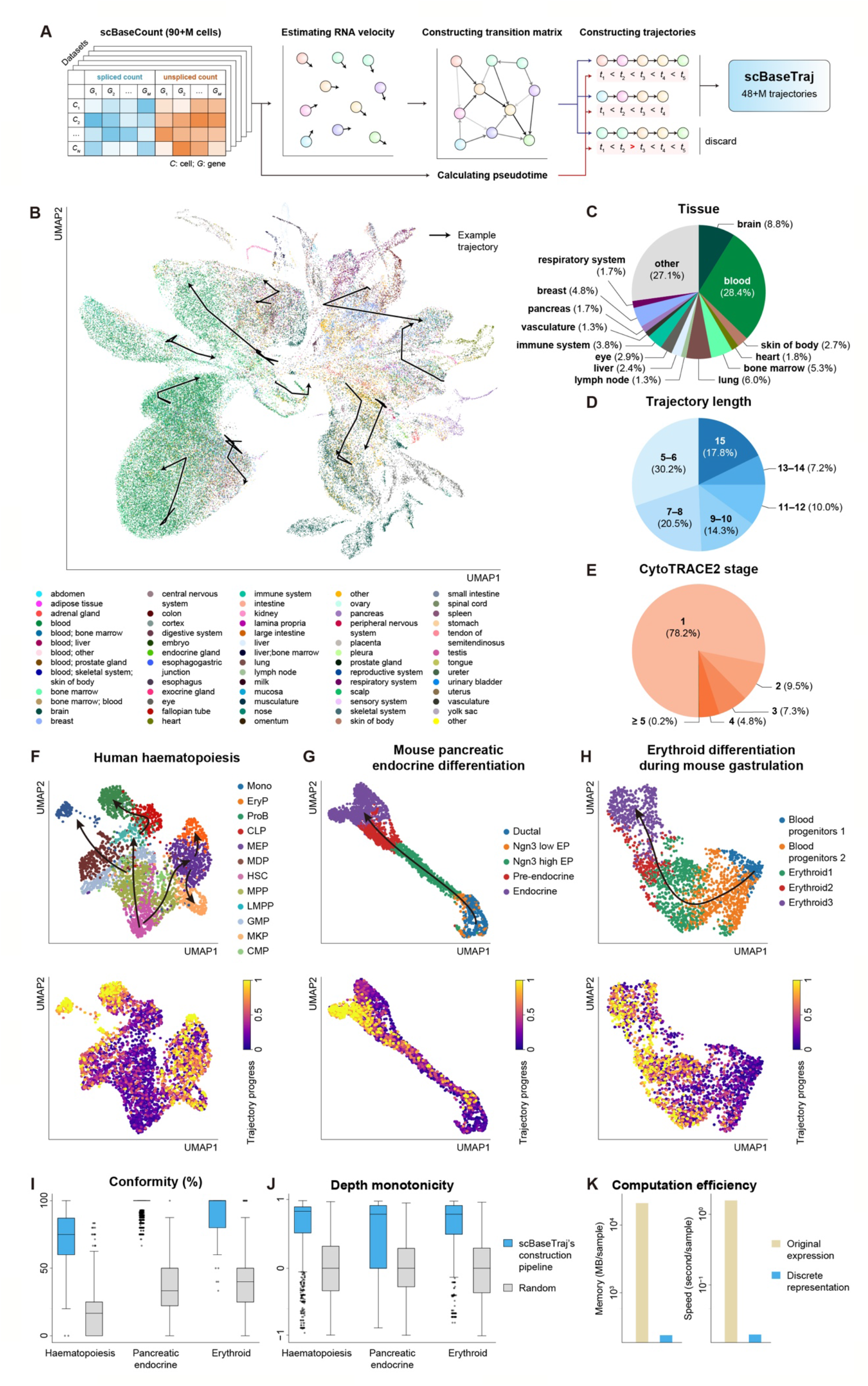
Construction and characterization of scBaseTraj. (A) Construction process of scBaseTraj. We first estimate RNA velocity from spliced and unspliced single-cell data and derive cell–cell transition probability matrix. This matrix is then used to construct multi-step cellular trajectories, constrained by transcriptome-derived pseudotime. (B) UMAP illustration of scBaseTraj with representative example trajectories overlaid. Cells are colored by tissue origins, with cells along a representative trajectory connected by lines. (C) Tissue composition of scBaseTraj. (D) Distribution of trajectory lengths in scBaseTraj. The frequency generally decreases with increasing length, except for length 15, which reflects the preset maximum trajectory length. (E) Distribution of CytoTRACE2 stages across all trajectories in scBaseTraj. (F– H) Visualizations of external datasets with well-characterized lineage hierarchies: human hematopoiesis (F), mouse pancreatic endocrine differentiation (G), and mouse erythroid differentiation (H). For each subfigure, the top panel shows the UMAP colored by cell types with arrows indicating differentiation direction, and the bottom panel shows the UMAP colored by trajectory progression in each generated trajectory. The trajectory progress is the normalized time step of the cell in the trajectory. For cells appearing in multiple trajectories, the maximum time step is visualized. (I–J) Conformity score (I) and depth monotonicity correlation (J) comparing the scBaseTraj’s construction with the random sampling pipeline in each dataset. (K) Computational efficiency comparison between models operating in original expression space and discrete representation space.

Starting from over 90 million single-cells with spliced and unspliced transcript counts derived from scBaseCount^8^, we inferred temporally ordered cellular trajectories by integrating velocity-informed state transitions with biologically informed directionality constraints. RNA velocity was estimated using scVelo^17^, and velocity-informed transition probabilities between cells were derived with CellRank2^18^ to assemble multi-step trajectories. These trajectories were further constrained and quality-controlled using pseudotime calculated by CytoTRACE2^19^ to ensure biologically plausible cell state transitions (Methods). Through this procedure, static single-cell snapshots were converted into explicit temporal sequences suitable for the autoregressive forecasting task.

The resulting dataset comprises more than 48 million trajectories spanning 71 tissues, with each trajectory covering an average of 9.3 consecutive states (Figure 2B–D). Approximately three-quarters of trajectories are confined within a single CytoTRACE2 stage, corresponding to relatively stable cell states, while the remaining trajectories span multiple CytoTRACE2 stages and capture dynamic state transitions (Figure 2E). Both types of trajectories were retained during training, enabling the model to learn representations of stable cell states as well as processes of dynamic state change.

Because scBaseCount does not include cell-level annotations, we evaluated the biological plausibility of the constructed trajectories using external datasets with well-characterized lineage hierarchies (Figure 2F–H). In a hematopoietic differentiation dataset^20^ with established lineage structure^21,22^, we used the above-mentioned pipeline to construct cell trajectories, and colored cells in the original UMAP with their normalized time step in each trajectory. As shown in Figure 2F, trajectories inferred by the construction pipeline showed strong concordance with known lineage directions and depth ordering, with a mean conformity score of 74.8%, which assesses the percentage of trajectory steps aligned with the correct differential direction, and a mean depth monotonicity correlation of 0.63, which assesses whether a trajectory progresses coherently along the lineage hierarchy (Figure 2I–J). In contrast, randomly selected cells from the dataset and constructed trajectories (Methods) achieved a mean conformity score of only 19.0% and a near-zero depth monotonicity correlation (−0.003). Similar consistency was observed when applying the same construction pipeline to mouse pancreatic endocrine differntiation^23^ and erythroid differentiation during mouse gastrulation^24^ datasets (Figure 2G–J). These results indicate that the scBaseTraj construction pipeline reliably captures biologically meaningful temporal progression and suggest that scBaseTraj can provide reliable supervision for training forecasting models.

Using scBaseTraj as explicit temporal supervision, we trained CellTempo using the aforementioned two-stage architecture, comprising over 49.64 million parameters, to learn multi-step cellular dynamics by forecasting future states along inferred trajectories (Methods). CellTempo has high scalability and computational efficiency (Figure 2K and Figure S2), and can be trained on a single NVIDIA A100 GPUs with 80 GB of memory.

### Forecasting long-range lineage progression from single initial states

To evaluate the temporal generative capability of CellTempo, we assessed its ability to forecast multi-step cellular trajectories starting from a single observed state. For each trajectory in the test set, CellTempo was provided with the first time point and used to autoregressively generate the subsequent five time points.

We evaluated trajectory generation at four complementary levels: local transcriptional dynamics, long-range temporal progression, global state-transition topology, and cell-type and gene-program dynamics. To assess local transcriptional dynamics, we evaluated how well CellTempo captures temporal expression changes along real trajectories. Specifically, for each pair of adjacent time points in the ground-truth trajectories, we identified the 500 genes exhibiting the largest expression changes and computed their change vectors. We then quantified the concordance between predicted and real expression changes for these genes using Pearson correlation coefficient (PCC) and Spearman correlation coefficient (SCC) (Methods). As shown in Figure 3A, CellTempo achieved consistently high PCC and SCC values, indicating that it accurately recapitulates dominant transcriptional dynamics underlying cell state transitions. As expected, correlations gradually decreased at later prediction steps, reflecting the accumulation of uncertainty in multi-step forecasting. Nevertheless, CellTempo maintained substantial agreement with real trajectories across all time points, demonstrating its ability to generate biologically coherent future cell states and preserve key dynamic trends starting from a single initial condition.

**Figure 3.**
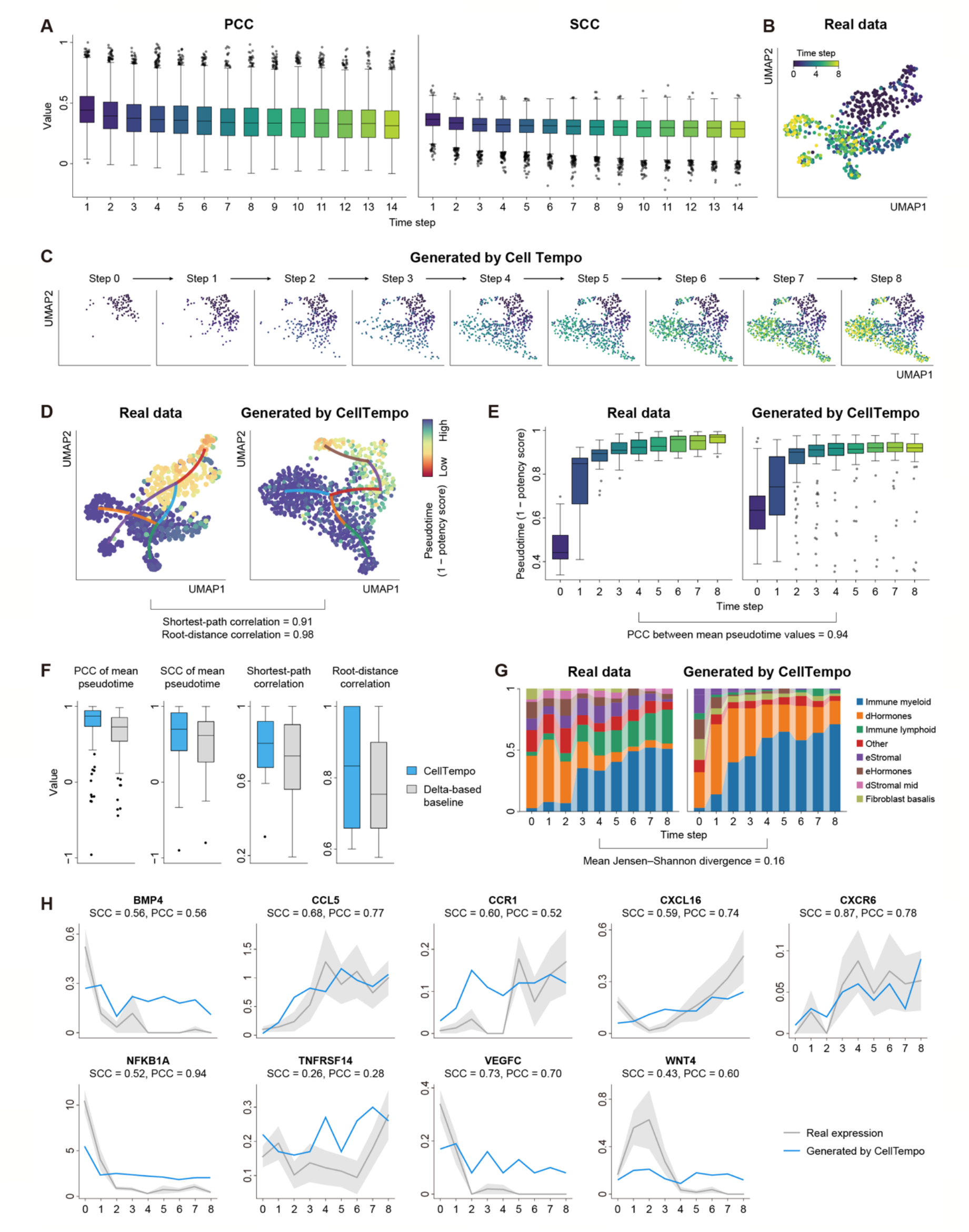
Evaluation of CellTempo-generated trajectories. Unless otherwise stated, panels (B– E, G–H) are based on the human uterus dataset. (A) Performance of CellTempo in predicting the 500 most changed genes in each test-set trajectory. Because each trajectory contains a maximum of 15 steps, we predicted the next 14 future steps beyond the initial cell. (B) UMAP of cells in real trajectories, colored by time step. (C) UMAP of cells at each step along the CellTempo-generated trajectories, colored by time step. (D) UMAP of cells in the real trajectories and generated trajectories colored by pseudotime, defined as 1 – CytoTRACE2 potency score. Lines represent high-probability transitions between clusters based on a PAGA-derived cluster transition graph. (E) Pseudotime of real and generated cells at each time step. (F) Performance of CellTempo-generated trajectories across all test datasets compared with the delta-based extrapolation baseline. (G) Cell-type composition at each time step in real and generated trajectories. (H) Temporal dynamics of marker gene expression across time steps .

To assess long-range temporal progression, we evaluated CellTempo across all test datasets individually. For each dataset, we computed pseudotime using CytoTRACE2 and selected the 100 cells with the lowest pseudotime as starting points. From each starting cell, we constructed real trajectories of length nine and, in parallel, generated trajectories of the same length using CellTempo.

Using a human uterus dataset^25^ in the test set as an illustrative example, we visualized both real and generated trajectories with UMAP (Figure 3B–C). The generated trajectories exhibit clear temporal ordering and progression patterns that closely resemble those of real trajectories. We then computed pseudotime independently on both trajectories (Figure 3D). Step-wise pseudotime distributions showed highly concordant trends (Figure 3E), with a PCC of 0.94 between real and generated mean pseudotime values (Methods).

As a simple temporal baseline, we constructed a delta-based extrapolation strategy that predicts the next time point by adding the average expression change at each corresponding step, estimated from training trajectories of the same tissue, to the current state. Across all test datasets, PCCs and SCCs between real and CellTempo-generated mean pseudotime reached 0.74 ± 0.35 (mean ± sd) and 0.62 ± 0.36, compared with 0.63 ± 0.33 and 0.54 ± 0.34 for the delta-based baseline (Figure 3F), indicating that CellTempo more faithfully captures temporal progression by modeling condition-dependent state transitions rather than relying on population-level average dynamics.

Beyond continuous progression, we evaluated whether CellTempo preserves global state-transition topology. We applied PAGA^11^ to infer cluster-level transition graphs from both real and generated trajectories (lines in Figure 3D). Structural similarity was quantified using shortest-path correlation and root-distance correlation (Methods). Across all test datasets, CellTempo achieved values of 0.78 ± 0.17 and 0.84 ± 0.16 (mean ± sd) for these metrics, respectively, outperforming the delta-based baseline, which reached 0.71 ± 0.22 and 0.77 ± 0.13 (Figure 3F). These results indicate that CellTempo more accurately recovers branching structure and global lineage topology.

We further examined cell-type composition dynamics in the uterus dataset using CellTypist^26^ (Figure 3G, Methods). Both real and generated trajectories show highly concordant temporal trends, with immune myeloid cells gradually increasing and hormone-related cells decreasing over time. The mean Jensen–Shannon divergence between real and generated cell-type compositions across steps is 0.16, indicating a high degree of similarity. Consistent results were observed at the gene level: temporal expression patterns of known marker genes^25^ were well preserved in generated trajectories, showing strong concordance with real trajectories (mean Pearson correlation = 0.65, Figure 3H). Similar patterns were consistently observed in an additional test dataset of human immune cells (Figure S3).

Together, these results demonstrate that CellTempo can generate realistic, long-range cellular trajectories from a single observed state, capturing continuous progression, branching topology, and coordinated cell-type and gene-expression dynamics.

### Reconstructing cellular potential landscapes and exploring cell-fate redirections

A fundamental question in cell biology is how cells traverse different states in processes such as differentiation, lineage commitment, regeneration, and disease progression. Waddington’s epigenetic landscape offers a conceptual framework for illustrating such processes, in which cells reside at the top of a rugged potential landscape and progressively differentiate by descending into lineage-specific valleys^27,28^. Recovering such a potential landscape directly from single-cell data and predicting how cells may transition on this landscape is crucial for understanding the biological process^29^. Because CellTempo is trained autoregressively to forecast future cell states along cellular trajectories, its predictive distribution naturally encodes both directional tendencies and uncertainty in cell state transitions. We leveraged this property to construct a data-driven cellular potential landscape grounded in learned temporal dynamics and to simulate cell state transitions on this landscape (Figure 4A).

**Figure 4.**
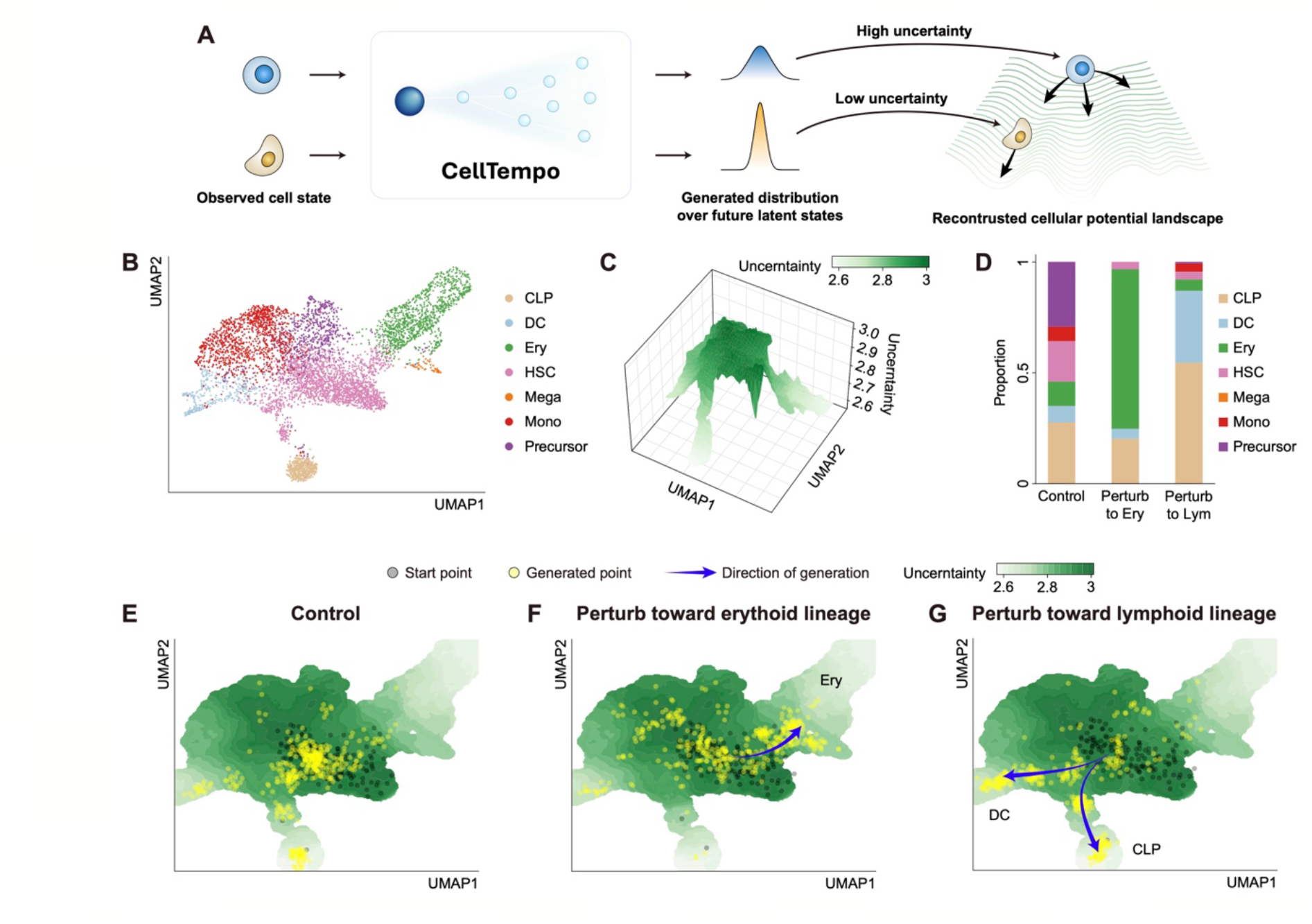
Reconstruction of the cellular potential landscape with CellTempo. (A) Schematic of cellular potential landscape reconstruction. Starting from observed cells in a given dataset, CellTempo autoregressively generates possible future cell states. Generated-cells with concentrated distributions are regarded as cells with low uncertainty, and *vice versa*. The uncertainty is used to reconstruct the cellular potential landscape. (B) The UMAP of the bone-marrow hematopoiesis dataset colored by cell type. (C) Overview of the reconstructed potential landscape colored by uncertainty score. (D) Cell-type composition of generated cellular trajectories under different perturbation conditions. (E–G) Visualizations in the potential landscape of generated cells (yellow) starting from HSCs (gray) without perturbation (E), under genetic perturbations toward erythroid lineage (F), and under genetic perturbations toward lymphoid lineage (G).

Specifically, we defined an uncertainty score for each cell in the specific dataset to measure its state transition potential. At each forecasting step, CellTempo outputs a probability distribution over all possible future latent states. The concentration of this distribution reflects the degree of determinism in a cell’s transition, whereas broader distributions indicate greater uncertainty and plasticity. To quantify the uncertainty, we computed the entropy of the predicted probability distribution for each of the latent tokens representing a cell, and defined a cell-level uncertainty score as the average entropy across all tokens in its discrete representation (Methods). Low uncertainty corresponds to strongly directed transitions, analogous to cells residing in deep potential basins, whereas high uncertainty indicates greater plasticity and multiple possible future fates. By projecting these entropy-derived uncertainty scores onto a two-dimensional UMAP embedding, we obtained a continuous, model-inferred potential landscape.

Here, we employed a human bone-marrow hematopoiesis dataset^30^ to conduct potential landscape reconstruction experiments. This dataset encompasses highly pluripotent hematopoietic stem cells (HSCs) alongside more differentiated lymphoid cells and erythroid cells. The inferred potential landscape by CellTempo exhibits a biologically coherent organization (Figure 4B–C). HSCs occupy regions of high inferred potential, consistent with their multipotent state, while progressively differentiated populations such as common lymphoid progenitors (CLPs), erythroid progenitors, and granulocyte–monocyte lineages localize to lower regions of the landscape. Smooth transitions from high- to low-potential regions follow known lineage branching patterns, with cells gradually diverging toward distinct fates, indicating that CellTempo successfully captures the intrinsic directional tendencies of hematopoietic differentiation.

Unlike pseudotime methods that impose one-dimensional orderings, or trajectory-inference approaches that connect observed states without modeling uncertainty, the CellTempo-derived potential landscape forms a continuous energy-like surface informed by probabilistic state-transition dynamics. This representation supports not only retrospective interpretation of cellular potential, but also prospective simulation of how perturbations on intrinsic drivers reshape fate decisions. As an example, we selected HSCs as starting points and generated cell-fate trajectories under both unperturbed and perturbed conditions (Methods). To lead differentiation toward the erythroid lineage, we up-regulated key erythroid transcription factor (TF) modules (GATA1, KLF1, GFI1B, NFE2, ZFPM1)^31-35^ while suppressing modules associated with lymphoid specification (SPI1, IKZF1/3, TCF3, EBF1, FLT3, IL7R)^36-41^. Under these genetic perturbations, CellTempo generated trajectories with a markedly increased fraction of erythroid-fated cells compared with the unperturbed condition. Conversely, perturbations promoting lymphoid-associated gene programs redirected differentiation toward lymphoid progenitors (Figure 4D). These perturbation-induced fate shifts are reflected directly on the potential landscape. When generated cells are overlaid onto the landscape, erythroid-promoting perturbations funnel cells preferentially into the erythroid basin, whereas lymphoid-promoting perturbations redirect trajectories toward the lymphoid branch (Figure 4E–G).

Together, these results demonstrate that CellTempo learns a coherent and biologically meaningful potential landscape over cell state space, capturing intrinsic differentiation potential and responding predictably to perturbations of driver gene modules. By unifying autoregressive forecasting with landscape reconstruction, CellTempo provides a principled framework for studying how dynamic gene-regulatory programs and external perturbations shape cell fate decisions.

### Forecasting long-term induction outcomes for iPSC-derived OPC fate engineering

Directed differentiation from induced pluripotent stem cells (iPSCs) provides a powerful system for generating therapeutically relevant cell types, but identifying effective induction strategies remains largely empirical^42,43^. Oligodendrocyte precursor cells (OPCs) are an important target of such efforts because they give rise to myelinating oligodendrocytes in the central nervous system and provide a cellular basis for potentially treating demyelinating and neurodegenerative disorders^44^. However, efficient OPC generation from iPSCs remains challenging, as oligodendrocyte lineage specification requires coordinated developmental transitions rather than a simple immediate transcriptional response ^44^. We therefore asked whether CellTempo could be used to prioritize OPC differentiation strategies by forecasting their long-term consequences on cell-fate trajectories.

We applied CellTempo to a published longitudinal single-cell RNA-sequencing dataset tracking the differentiation of human iPSCs into mixed populations of neurons, astrocytes, and oligodendrocyte-lineage cells ^45^. We focused on the day-30 proliferative glial progenitor cells (GPCs), which can differentiate into astrocytes or OPCs, making it a suitable starting population for forecasting perturbation-induced OPC formation.

We first investigated whether CellTempo could predict long-term fate changes following perturbation of regulatory genes associated with OPC differentiation. We curated a set of genes implicated in oligodendrocyte specification and up-regulated their expression in day-30 proliferative GPCs (Methods). The perturbed and unperturbed cells were then used as starting states for trajectory forecasting by CellTempo. Cell identities at each generated time step were assigned using marker-based annotation, and the abundance of OPC-lineage states, including pre-OPCs and OPCs, was monitored throughout the predicted trajectories. Compared with the unperturbed condition, up-regulation of OPC-associated regulatory genes substantially increased the abundance of OPC-lineage cells along the predicted trajectories (Figure 5A), demonstrating that CellTempo can capture the long-term cell-fate consequences of changes in key developmental regulatory programs.

**Figure 5.**
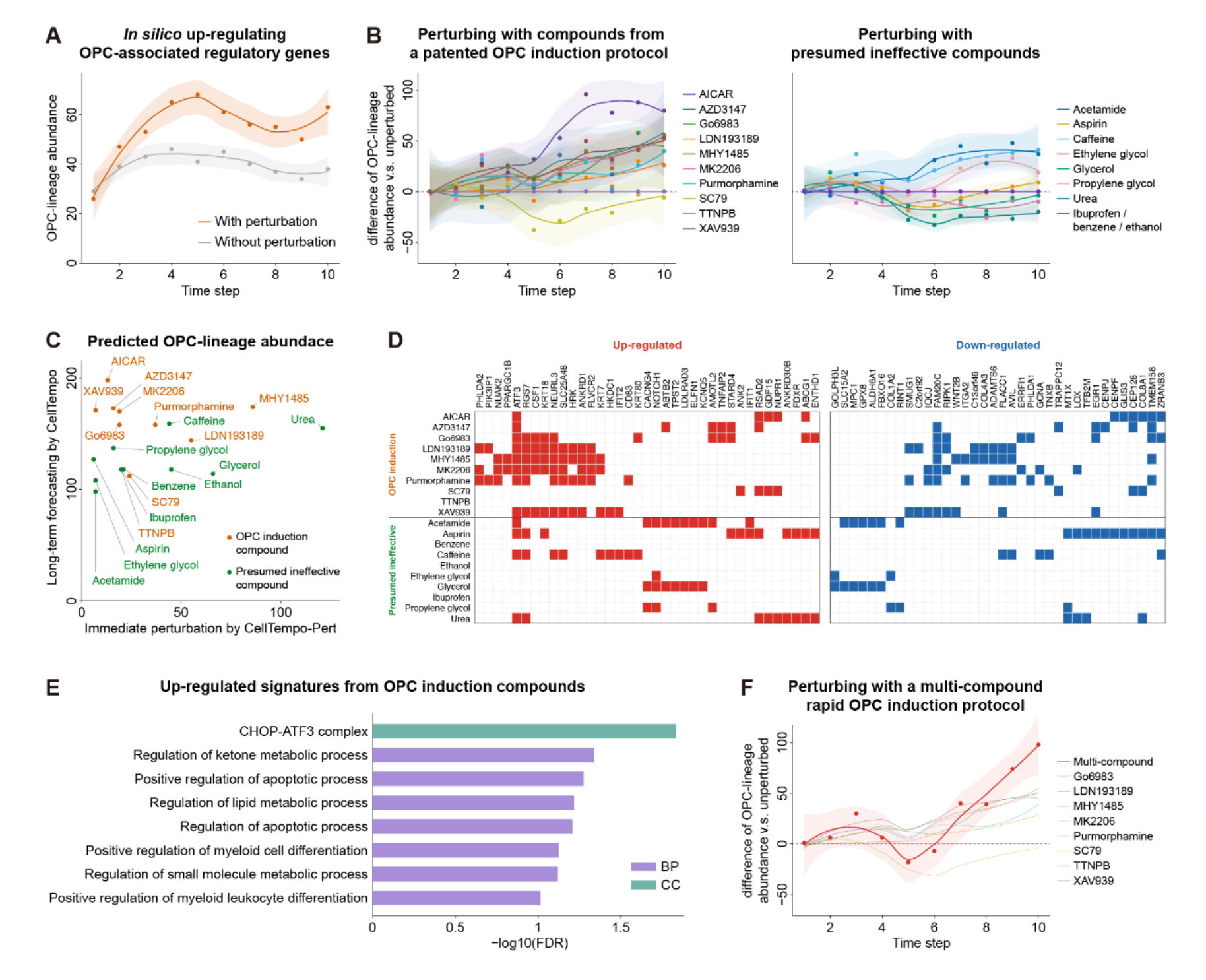
Forecasting OPC-directed differentiation from iPSC-derived day-30 proliferative GPCs with CellTempo. (A) Long-term trajectory forecasting after *in silico* up-regulation of OPC-associated regulatory genes with CellTempo. (B) Prediction of long-term effects of individual compounds on OPC induction with CellTempo. Curves represent loess-smoothed trajectories, with shaded areas indicating 95% confidence intervals. Values represent changes in OPC-lineage abundance relative to the unperturbed trajectory. (C) Comparison between immediate perturbation responses predicted by CellTempo-Pert and long-term fate outcomes predicted by CellTempo. (D) Compound-specific perturbation signature genes identified relative to DMSO-predicted states for each compound. Only recurrently perturbed genes across compounds are shown. (E) Gene Ontology (GO) enrichment analysis of perturbation signatures. Up-regulated and down-regulated signatures from OPC induction compounds, as well as signatures from presumed ineffective compounds, were analyzed separately. Only terms with FDR < 0.1 were shown. No significant enrichment was detected for down-regulated signatures from OPC induction compounds or either direction of signatures from presumed ineffective compounds. CC, cellular component; BP, biological process. (F) Prediction of long-term effects of a multi-compound OPC induction protocol with CellTempo. Individual compound effects within the protocol are shown as thin lines. Curves represent loess-smoothed trajectories, with shaded areas indicating 95% confidence intervals. Values represent changes in OPC-lineage abundance relative to the unperturbed trajectory.

We next asked whether CellTempo could be used to evaluate the long-term effects of individual compounds used in chemical OPC induction protocols. To model chemical perturbations, we extended the CellTempo architecture to incorporate drug information as an explicit conditional input. Specifically, we formulated drug response as a two-step transition, where untreated control cells define the initial state and drug-treated cells represent the immediate perturbed state. Molecular representations of compounds, computed using UniMol2^46^, together with contextual features including cell line and dosage, were encoded as conditioning tokens and appended to the cellular token sequence. We then fine-tuned CellTempo on the Tahoe-100M perturbation dataset^47^ to predict the discrete cellular representation following drug exposure, generating CellTempo-Pert, a perturbation-conditioned variant of the model (Figure S4, Methods).

We selected several compounds from a patented rapid iPSC-to-OPC differentiation strategy^48^, together with several common compounds that we expected to have little or no effect on OPC differentiation (Methods). For each compound, we first used CellTempo-Pert to predict the immediate transcriptional responses of day-30 proliferative GPCs to both the compound and DMSO. Using the DMSO-predicted state as a reference helped reduce transcriptional changes introduced by the perturbation model itself and highlight responses specific to each compound. The top 12 genes up- or down-regulated in the compound-predicted state relative to the DMSO-predicted state were then applied as *in silico* perturbations to the same starting population. We subsequently used CellTempo to forecast the long-term differentiation trajectories of these perturbed cells.

After long-term forecasting, most compounds selected from the OPC induction protocol generally produced sustained increases in the abundance of OPC-lineage cells, whereas the presumed ineffective compounds showed little or no enhancement of OPC differentiation (Figure 5B). Notably, SC79, a compound from the induction protocol, decreased the predicted OPC-lineage abundance compared with the control condition. This apparently counterintuitive effect is consistent with the stage-dependent behavior of SC79 reported in the patent. In the early OPC induction stage, SC79 was included as an AKT pathway activator in the optimized formulation. However, in a later SOX10-focused optimization model, SC79 showed the strongest negative contribution among the tested factors, suggesting that the effect of AKT activation depends on the developmental stage. Because our analysis started from proliferative GPCs rather than pluripotent cells, these results suggest that while AKT activation may facilitate early OPC specification, its continued activity at later progenitor stages may negatively affect progression toward a more mature OPC state.

We further compared the long-term outcomes predicted by CellTempo with the immediate responses predicted by CellTempo-Pert (Figure 5C). Interestingly, several compounds, such as glycerol, and urea, generated appreciable numbers of OPC-lineage cells in the immediate predictions, as inferred by marker-based cell type annotation. However, these apparent OPC-like states were not maintained during subsequent trajectory forecasting by CellTempo, suggesting that transient transcriptional responses induced by perturbations may mimic OPC marker expression without representing genuine progression toward the OPC fate. In contrast, XAV939, AICAR, and Go6983, which are components of OPC induction protocols, produced relatively few OPC-lineage cells in the immediate CellTempo-Pert predictions but showed a gradual and sustained increase in OPC abundance over future prediction steps. These contrasting temporal patterns demonstrate that immediate perturbation responses alone can be insufficient, and potentially misleading, for evaluating the effects of compounds on cell fate. By extending predictions beyond the initial perturbation state, CellTempo distinguished transient OPC-like transcriptional signatures from perturbations that drive persistent progression along the OPC differentiation trajectory, thereby enabling more reliable prioritization of compounds with long-term OPC-promoting potential.

To investigate why immediate perturbation predictions did not reliably reflect long-term outcomes, we examined the compound-specific perturbation signatures used for long-term trajectory forecasting. These signatures consisted of the top 12 genes up- or down-regulated relative to the DMSO-predicted state for each compound. Across compounds included in the OPC induction protocols, several genes, such as ATF3, RGS7, and NEURL3, were recurrently perturbed (Figure 5D). Functional enrichment analysis of these signature genes revealed significant associations with differentiation-related and apoptosis-related processes (Figure 5E), suggesting the activation of coordinated biological programs associated with fate progression. In contrast, perturbation signatures generated by presumed ineffective compounds showed limited recurrence across compounds and lacked significant functional enrichment.

These results suggest that immediate perturbation responses may not be sufficient to predict long-term cell-fate outcomes. Although some compounds induced transcriptional changes that were sufficient to generate OPC-lineage-like cells in the immediate prediction, these responses did not necessarily represent activation of the regulatory programs required for continued OPC differentiation. Instead, they may reflect transient or partial transcriptional responses to compound exposure that mimic aspects of the OPC state without driving stable fate progression. Consequently, these cells failed to develop into sustained increases in OPC-lineage abundance during long-term CellTempo forecasting. In contrast, compounds that induced recurrent and coordinated differentiation-associated programs showed progressive promotion of OPC-lineage formation. These results demonstrate that CellTempo can distinguish transient perturbation responses from molecular changes that are ultimately translated into long-term cell-fate transitions.

Finally, we asked whether CellTempo could extend from evaluating individual compounds to assessing multi-compound OPC induction strategies. We evaluated a multi-compound rapid OPC induction protocol described in the published patent^48^. CellTempo-Pert was first used to predict the immediate transcriptional responses of day-30 proliferative GPCs to each constituent compound and to DMSO. Differentially expressed genes relative to the DMSO-predicted state were then combined across all compounds to construct an aggregate perturbation signature representing the protocol-level molecular effect (Methods). This signature was applied to the starting population, and CellTempo was used to forecast the resulting long-term differentiation trajectories. The multi-compound combination produced larger sustained increases in OPC-like cell abundance than all its individual constituent compounds, suggesting that CellTempo can integrate the effects of multiple perturbations at the protocol level (Figure 5F).

Together, these results demonstrate that CellTempo can move beyond immediate perturbation responses to distinguish transient effects from sustained cell-fate changes, predict the long-term consequences of multi-compound induction strategies, and facilitate the rational design and optimization of directed differentiation protocols.

## Discussion

In this study, we introduce CellTempo, an autoregressive generative model for forecasting long-range cell state transitions from static single-cell transcriptomes. Pretrained on the large-scale transition database scBaseTraj, CellTempo learns recurrent patterns of cell state progression and maps a current state to a distribution of plausible future states. Across diverse evaluations, CellTempo accurately forecasts multi-step transcriptional trajectories, reconstructs coherent differentiation structure, infers cellular potential landscapes, and predicts long-term cellular responses to perturbations. These findings demonstrate that long-range cellular dynamics can be inferred from snapshot measurements, expanding the scope of what static single-cell data can reveal.

At a conceptual level, CellTempo treats cell state transitions as a predictive dynamical process that can be forecast, perturbed, and simulated, rather than merely inferred *post hoc*. In this perspective, forecasting trajectories corresponds to generating possible future states, potential landscapes emerge from the distribution of these futures, and perturbation effects are represented as shifts in predicted trajectories. By linking current states to probabilistic futures, CellTempo provides a predictive view of cellular behavior and illustrates how snapshot measurements can be interpreted in a dynamical context. Thus, CellTempo represents a step toward a “world model” of cells, in the sense that it learns probabilistic rules governing cell state transitions over time rather than static representations.

A central advance of CellTempo is its extension of perturbation modeling from immediate response prediction to long-term outcome forecasting. Most existing *in silico* perturbation models estimate transcriptional states shortly after an intervention^49-51^, yet such responses can be transient and may not reflect long-term changes in cell fate. CellTempo addresses this gap by propagating predicted perturbation responses through future state space to determine their downstream consequences. Applied to OPC cell-fate engineering, CellTempo prioritized interventions according to whether their immediate responses ultimately led to sustained OPC formation. The resulting long-term ranking differed from that based on immediate responses alone and further enabled the evaluation of multi-compound induction strategies. These results identify temporal horizon as a fundamental dimension of perturbation modeling and show that perturbations should be evaluated by their long-term cellular consequences rather than by short-term transcriptional responses alone.

There are several important opportunities to further extend the CellTempo model. First, the temporal supervision used to train CellTempo is derived from computationally inferred trajectories rather than direct longitudinal measurements. Although constrained by RNA velocity and pseudotime, these trajectories inevitably reflect assumptions of the underlying inference methods. Integrating additional sources of temporal information, such as lineage tracing experiments^52^, metabolic labeling^53^, or emerging technologies that repeatedly profile the same cell over time^54^ could provide more direct supervision and further improve model fidelity. Second, CellTempo currently operates on scRNA-seq data alone, capturing only one layer of cell state. Extending the model to incorporate multimodal measurements, including chromatin accessibility, protein abundance, and spatial context, represents an important step toward a more comprehensive and mechanistic representation of cellular dynamics. In addition, as CellTempo encodes cell states through a VQ-VAE–based discrete representation that emphasizes global state configurations, genetic perturbations are more naturally modeled as coordinated gene-module–level changes rather than isolated single-gene interventions. This design choice limits direct gene-level perturbation resolution but reflects the model’s focus on state-level dynamics. In this context, our chemical perturbation experiments provide an important proof of principle, demonstrating that perturbation-aware fine-tuning enables CellTempo to capture immediate responses and propagate their effects forward in time, suggesting a modular strategy in which gene-level perturbation models predict initial perturbed states, while CellTempo focuses on forecasting downstream temporal state evolution. Finally, while autoregressive modeling combined with VQ-VAE discretization offers a flexible and scalable solution for temporal generation, alternative generative paradigms, such as diffusion-based models^55-57^, may provide complementary advantages for modeling complex, high-dimensional biological systems.

Looking forward, CellTempo provides a foundation for studying cellular dynamics across development, disease progression, and therapeutic response. By learning how cells change rather than only how they appear, CellTempo enables a new class of *in silico* experiments in which lineage progression, cell fate decision, and drug responses can be explored and designed within a unified generative model. We anticipate that the datasets, modeling paradigm, and analyses presented here will support future efforts toward predictive and mechanistic models of cell fate evolution.

## Methods

### Architecture of CellTempo

The model of CellTempo consists of two major components: a cell-level vector-quantized variational autoencoder (VQ-VAE) that maps continuous gene-expression profiles into a discrete latent space, and a Transformer-based autoregressive model that generates sequences of discrete cell representations to model temporal trajectories. These two components enable scalable generation of long-range cell state evolutions.

#### Cell VQ-VAE for learning discrete cellular representations

The cell VQ-VAE, consisting of an encoder, a vector-quantization codebook, and a decoder, is used to encode the cell expression vector into discrete latent representations and decode the representation back to expression data. Let each cell be represented by a raw count vector **x** ∈ ℝ^*n*^, where *n* is the number of genes. The encoder *E*(⋅) maps the gene-expression vector into a low-dimensional continuous embedding:

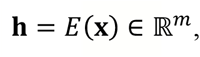

where *m* is the encoder output dimension. We reshape **h** into a sequence of *i* latent codes, each of dimension *j*, such that *m* = *i* × *j*:

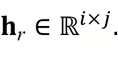

The codebook is a learnable set of *K* embedding vectors:

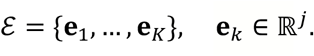

For each latent code **h**_*t*_ in **h**_*r*_ (*t* = 1, …, *i*), we replace it with its nearest neighbor in the codebook:

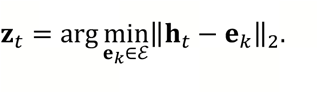

Let *c*_*t*_ ∈ {1, …, *K*} denote the index of the selected codebook vector. The discrete representation of a cell is then given by the index sequence:

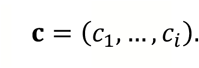

To reconstruct the expression profile, we map **c** back to their corresponding codebook vectors:

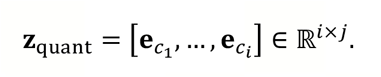

The decoder *D*(⋅) transforms this representation into the mean parameter of a Negative Binomial (NB) distribution for each gene:

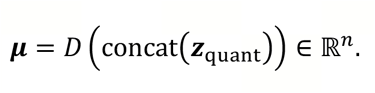

Each gene additionally has a learnable NB dispersion parameter ***θ*** ∈ ℝ^*n*^, shared across all cells in the dataset. The reconstructed counts are sampled from:

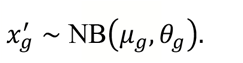

The VQ-VAE is trained by minimizing the standard loss:

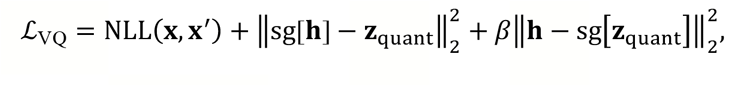

where NLL is the negative log-likelihood, sg[⋅] denotes the stop-gradient operator, and *β* controls the commitment loss. This VQ-VAE converts each cell into a fixed-length discrete token sequence, providing a compact and stable representation for downstream autoregressive modeling.

#### Transformer backbone for autoregressive generation of cell trajectories

We next train a Transformer model to generate cell representation autoregressively in the discrete latent space. For a trajectory containing *T* ordered cells, we encode each cell into a discrete index sequence. First, for the discrete token sequence of a cell **c**^(*t*)^, it is augmented with start and end markers:

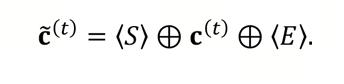

Cells are then concatenated into a trajectory sentence:

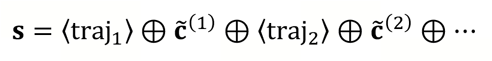

The final token vocabulary consists of codebook entries, trajectory-index tokens (⟨traj_1_, … ⟩), and special tokens (⟨*S*⟩, ⟨*E*⟩, ⟨pert⟩, ⟨control⟩, etc.), yielding a total vocabulary size:

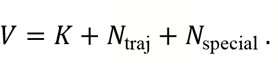

To encode temporal position along the trajectory, we introduce a cell-position token indicating which cell a token belongs to:

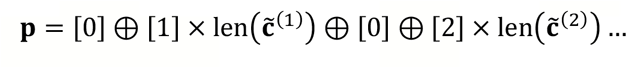

We use separate embedding layers to obtain the embedding of trajectory sentence and cell position token, and feed them into the Transformer model. We use the next token prediction method to train the transformer model. Let **s** = (*w*_1_, …, *w*_*L*_) denote the full trajectory token sequence. The Transformer receives **s** as input under causal masking, ensuring that 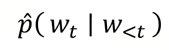 depends only on previous tokens. The training objective is the cross-entropy loss:

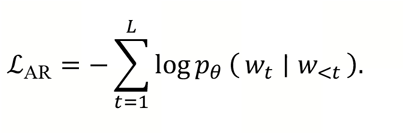

During inference, given an initial cell represented by **c**^(1)^, we construct the initial sequence:

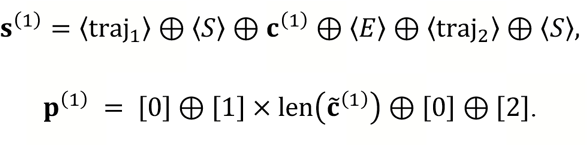

The Transformer then autoregressively samples the discrete representation of the next cell 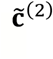. We then concatenate the new inter-cell tokens to the current **s** and construct:

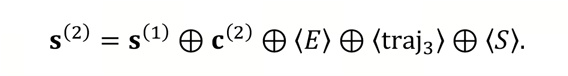

The cell position tokens are updated in parallel:

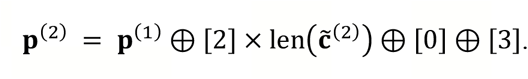

This procedure is repeated iteratively until the model generates a trajectory of the specified length. After that, each predicted cell can be decoded through the VQ-VAE decoder to obtain full gene-expression profiles.

This two-stage architecture (discrete representation learning via VQ-VAE followed by autoregressive sequence modeling) enables efficient and biologically coherent generation of long-range cellular trajectories. Discretization greatly reduces computational cost while retaining expressive power, and the Transformer backbone captures temporal dependencies that govern cell state transition.

### Model training and implementation details

Model training includes two separate stages: training the cell VQ-VAE model, and training the Transformer-based autoregressive model. All training is performed on four NVIDIA A100 GPUs (80 GB memory each). All inference experiments are performed on a single NVIDIA A100 GPU (80 GB).

The cell VQ-VAE consists of a three-layer multilayer perceptron (MLP) encoder, a three-layer MLP decoder, and a vector-quantization codebook containing 512 codes, each with a feature dimension of 24. Each cell is represented using 26 discrete codes. The VQ-VAE is trained for 7 × 10^5^ steps with a batch size of 64 per GPU (total batch size 256), using a learning rate of 1 × 10^−4^ with 500 warm-up steps.

The Transformer-based autoregressive model includes 12 layers and 12 attention heads, where each token is embedded into a 600-dimensional vector space. The model is trained for 1 × 10^4^ steps with a batch size of 32 per GPU (total batch size 128), a learning rate of 1 × 10^−4^, 1,000 warm-up steps, and a dropout rate of 0.1

As a baseline comparison, we train autoregressive models directly in the full gene-expression space without discrete representation learning. For each cell, gene-expression profiles are represented using the same set of 18,791 protein-coding genes as in CellTempo. Trajectories are modeled as short sequences of length 2, corresponding to next-step prediction along inferred trajectories. The autoregressive model architecture and training protocol follow the same settings as the CellTempo Transformer backbone, where applicable, except that predictions are made directly in the continuous gene-expression space rather than in the discrete latent space.

### Construction of the trajectory dataset scBaseTraj

To assemble a large-scale corpus of cellular trajectories for model pretraining, we construct trajectories independently for each dataset and then aggregate them into a trajectory dataset scBaseTraj. For each dataset, we integrate pseudotemporal ordering, RNA velocity, and probabilistic state-transition modeling to derive biologically coherent cell-state progression paths. The overall procedure consists of four major steps.

First, we compute pseudotime for all cells using CytoTRACE2^19^ (calculated as 1 − potency score), which provides a global ordering of cells based on transcriptional potential. Second, we estimate RNA velocity using scVelo^17^ to obtain directional information about short-term transcriptional dynamics. Third, we use CellRank2^18^ to integrate the velocity information into a cell–cell transition probability matrix, representing the likelihood of transitioning from one cell state to another. Finally, we construct trajectories by sampling paths through this transition matrix under biologically motivated constraints. Specifically, at each step, candidate next states are derived from the CellRank-derived matrix following the transition probabilities. To ensure directional and non-redundant progression, previously visited cells are excluded, preventing backtracking along the trajectory. To avoid local oscillations and redundant sampling of closely related states, immediate neighbors of the current cell are excluded from consideration. In addition, monotonic progression along developmental direction is enforced by removing candidate cells with lower CytoTRACE2 pseudotime than the current state. The next cell is sampled probabilistically in proportion to its transition probability from the remaining valid candidates, and the procedure is iterated until a full trajectory is constructed.

To reduce high-frequency noise and ensure sufficient temporal separation between steps, each trajectory is subsequently subsampled every three steps. This downsampling retains the general directional trends of cellular progression while discarding local fluctuations that would not meaningfully contribute to model training.

This standardized procedure is applied to 16,057 datasets from scBaseCount, and the resulting trajectories are aggregated to form the final large-scale trajectory dataset, scBaseTraj. One hundred datasets are randomly held out as an independent test set, with trajectories from the remaining datasets used for model training. To ensure consistency across datasets, gene identities are harmonized by restricting all analyses to 18,791 protein-coding genes.

### Validation of the scBaseTraj construction pipeline

#### Preprocess of human hematopoietic differentiation data

To evaluate the trajectory construction pipeline, we utilize the raw lineage tracing data provided by Weng et al.^20^, which contains single-cell data of human hematopoietic differentiation process. Since the official dataset does not contain spliced and unspliced information, we process the raw data before we construct the trajectories. Specifically, the data is preprocessed using the count function of the CellRanger-ARC^58^ pipeline. Subsequently, we employ the run10x function of Velocyto^10^ to convert the BAM format files into 10X-formatted files containing spliced and unspliced matrices. These spliced and unspliced data are subsequently used to calculate RNA velocity and construct cellular trajectories using the procedure described above.

#### Random construction baseline

To provide a baseline for our trajectory construction method, we generate trajectories using a random sampling strategy. Specifically, we first calculate the average trajectory length *N* obtained by our construction method on the test dataset, and then randomly sample *N* cells from the test dataset and arrange them in the order of sampling to form a trajectory. Repeating this sampling-and-ordering procedure yields a set of distinct random trajectories.

#### Quantification of differential direction agreement

We quantify the agreement of differential direction using a conformity score. Given a trajectory *T* = (*t*_1_, *t*_2_, …, *t*_*L*_) consisting of cell type labels, each transition (*t*_*i*_ → *t*_*i*+1_) is categorized as forward, stay, or other. Forward means the transition move along a valid parent → child edge in the lineage tree. Stay means the trajectory remains in the same annotated cell type. Other is any other situation. Let *N*_forward_, *N*_stay_, and *N*_other_ denote the counts of each type among the total of *L* − 1 transitions. We define the conformity score as:

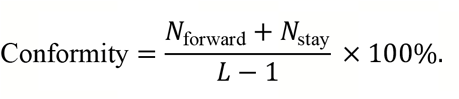

The score lies in the interval [0, 100%], where a score close to 100% indicates that the predicted trajectory consistently follows biologically valid developmental transitions, whereas a low score indicates frequent violations of lineage topology.

#### Depth monotonicity correlation

The depth monotonicity correlation aims to assess whether a trajectory progresses coherently along the developmental hierarchy. Each cell type *t*_*i*_ is assigned a differentiation depth *d*_*i*_ based on the lineage tree, where the root cell type has depth zero and depth increases along differentiation branches. For a trajectory *T* with depth sequence *D* = (*d*_1_, *d*_2_, …, *d*_*L*_), we compute the Pearson correlation coefficient (PCC) between the sequence and the temporal index of the trajectory steps (1,2, …, *L*). This metric quantifies the extent to which a trajectory follows a monotonic differentiation trend from progenitor toward more differentiated states. Values close to 1 indicate smooth and coherent progression along the developmental hierarchy, whereas values near 0 indicate the absence of a consistent directional pattern. Importantly, depth monotonicity correlation evaluates global developmental directionality across the entire trajectory, rather than the correctness of individual state transitions.

#### Comparing autoregressive training in latent space and full gene-expression space

To assess the biological fidelity between our two-stage strategy and the model trained directly in full gene-expression space, we evaluate cells generated by each approach using five complementary metrics: PCC, Spearman correlation coefficient (SCC), Local Inverse Simpson’s Index (LISI), Maximum Mean Discrepancy (MMD), and Area Under the Curve (AUC).

PCC and SCC measure linear and rank-based similarity between generated and real cells, respectively. For each metric, we compute the correlation for all generated–real cell pairs and reported their mean as the final score.

LISI quantifies the degree of mixing between real and generated cells in a shared neighborhood graph. We first construct a *k*-nearest neighbor (KNN) graph using 20 principal components (*k* = 10), and then compute LISI using the ilisi_graph function from the scIB package^59^.

MMD quantifies the distance between the distributions of real cells (*P*) and generated cells (*Q*) in a reproducing kernel Hilbert space (RKHS). Given kernel function *k*(⋅,⋅), the squared MMD is defined as:

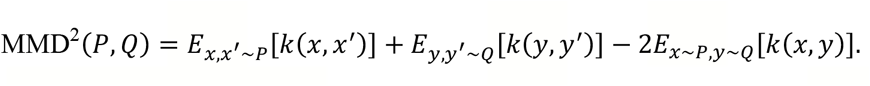

We compute MMD using the principal components of the real and generated cells.

The AUC score is derived from a Random Forest classifier trained to discriminate real from generated cells. Specifically, we train a Random Forest classifier with 1,000 trees and maximum depth of 5, using a random 75%/25% training–validation split. Performance was measured using the AUC under the receiver operating characteristic curve (ROC). An AUC close to 0.5 indicates that real and generated cells are indistinguishable, suggesting good biological realism, whereas an AUC closer to 1.0 indicates strong separability.

### Evaluation of generated cellular trajectories

#### Delta-based extrapolation baseline

We implement a delta-based extrapolation strategy as a non-autoregressive baseline for trajectory generation. This approach constructs trajectories by interatively applying average expression displacements estimated from real data.

Specifically, let 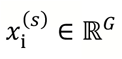 denote the gene expression vector of cell *i* at step *s*, where *G* is the number of genes. For each tissue *t*, we estimate the mean step-wise expression displacement from training trajectories as

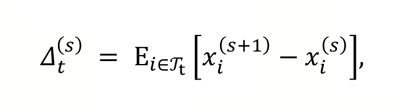

where *T*_t_ denotes the set of training trajectories from tissue *t*, and the expectation is taken over all cells observed at step *s* in these trajectories. Given a starting cell with expression *x*^(0)^, a trajectory of length nine is then generated by iterative propagation:

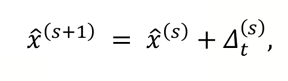

where the tissue *t* is determined by the dataset being evaluated. This strategy serves as a baseline for comparison with the trajectory generated by CellTempo.

#### Assessing local transcriptional dynamics

We evaluate how well CellTempo captures temporal expression changes along real trajectories. We first generate trajectories for cells in the scBaseTraj’s test set. For each pair of adjacent time points in the ground-truth trajectories, we identify 500 genes exhibiting the largest expression changes and computed their change vectors. We then quantify the concordance between predicted and real expression changes for these genes using PCC and SCC.

#### Assessing long-range temporal progression

For each dataset, we compute pseudotime using CytoTRACE2 and selected the 100 cells with the lowest pseudotime as starting points. From each starting cell, we construct real trajectories of length nine and, in parallel, generated trajectories of the same length using CellTempo. We then compute pseudotime independently on both trajectories. PCC is computed between real and generated mean pseudotime values. We do this in every test dataset, and the final PCC is obtained by averaging the PCC score of all test datasets.

#### Assessing global state-transition topology

To assess the similarity of global state-transition topology between real and generated trajectories, we compare PAGA-derived cluster-level transition graphs using two complementary metrics: shortest-path correlation and root-distance correlation.

To quantify global topological similarity, we compute a shortest-path correlation between the real and generated cluster graphs. Let *G*^(*r*)^ and *G*^(*g*)^ denote the cluster graphs derived from PAGA for the real and generated data, respectively. For each graph, we compute all-pairs shortest-path distances *d*^(*r*)^ and *d*^(g)^, and collect the finite entries of the upper-triangular distance matrix 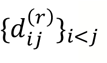 and 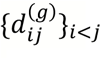. To make the comparison robust to graph size, we summarize each distance distribution by its empirical quantiles *Q*^(*r*)^ (*q*) and *Q*^(*g*)^ (*q*) for *q* ∈ [0.01,0.99] sampled on a regular grid. The shortest-path correlation is then defined as the SCC between the two quantile vectors. Values close to 1 indicate highly similar distributions of geodesic distances, reflecting comparable global graph shape and path-length structure, whereas lower or negative values indicate substantial topological divergence.

To evaluate whether the radial organization of the transition graphs is preserved, we assess similarity in distances from the inferred differentiation root using root-distance correlation. For each graph, the root cluster is defined as the cluster with minimal mean pseudotime, computed using diffusion pseudotime (DPT) in Scanpy, denoted *r*^(*r*)^ and *r*^(*g*)^ for the real and generated data, respectively. We then compute graph distances from the root to all nodes: 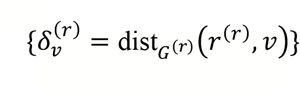 and 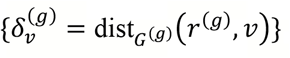. We compare the empirical quantile functions 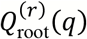 and 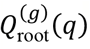 using SCC and define the result as the root-distance correlation. This metric captures how well the relative depth and radial expansion of branches from the inferred origin of differentiation are preserved in the generated trajectories, with higher values indicating closer agreement in global developmental organization.

#### Assessing cell-type dynamics

The cell-type composition is derived by CellTypist^26^. CellTypist is a supervised cell type annotation method that uses a logistic regression classifier trained on curated reference atlases to assign cell identities based on transcriptomic profiles. The difference between real and generated cell-type compositions is calculated by Jensen–Shannon divergence (JSD), a symmetric and smoothed measure of dissimilarity between two probability distributions. Given two discrete distributions *p* and *q* representing, for example, the cell-type composition of real and generated trajectories, JSD quantifies how different these distributions are while ensuring numerical stability and boundedness. Formally, JSD is defined as:

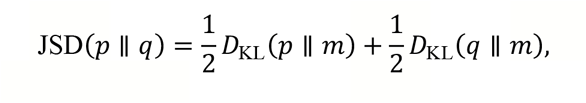

where 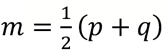 and *D*_KL_(*p* ∥ *m*) denotes the Kullback–Leibler divergence between *p* and *m*. To avoid numerical instability, all probabilities are clipped to a small positive constant before normalization.

### Construction of the cellular potential landscape

To construct a potential landscape that reflects the preferred directions and tendencies of cell state progression, we estimate a cell-level uncertainty score based on the autoregressive predictive behavior of the pretrained CellTempo model. For a given dataset, we treat every cell as a trajectory starting point and use the model to generate the next two future time steps.

During autoregressive generation, CellTempo outputs, for each token in the predicted cell state, a probability distribution over the discrete vocabulary. The concentration of this distribution reflects the model’s confidence in the predicted transition. For each generated cell state, we collect the logits for all tokens in its discrete representation and apply a softmax to obtain token-level probability distributions. Entropy is then computed as

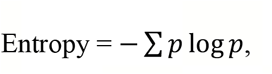

using a logits tensor *p* ∈ *R*^tokens×vocab_size^, with tokens denoting the number of tokens in the cell’s discrete representation and vocab_size the size of the model vocabulary.

The mean entropy across all tokens is used as the uncertainty score for that cell, and the uncertainty scores of the two generated future steps are averaged to produce a single developmental potential value for the starting cell. We then embed the original dataset into a two-dimensional UMAP space using the standard Scanpy preprocessing workflow and assign each cell’s uncertainty value as its *z*-axis coordinate. The resulting three-dimensional surface forms a landscape analogous to a Waddington potential landscape, with high-uncertainty cells residing at elevated regions and committed cells occupying lower-energy basins.

### Manipulating cell differentiation tendency on the potential landscape

To assess whether the learned cellular potential landscape captures biologically meaningful dynamic tendencies, we perform targeted perturbation experiments directly on the landscape by altering gene expression states of cells located in high-potential regions and examining the resulting changes in their predicted developmental trajectories. We first select a population of high-potential cells, such as hematopoietic stem cells (HSCs) in the bone marrow dataset, and use each as a starting point for generating three subsequent time steps with CellTempo.

In parallel, we generate perturbed versions of these starting cells by modifying gene expression levels in lineage-associated gene modules. For erythroid-directed perturbations, we up-regulate GATA1, KLF1, GFI1B, NFE2, and ZFPM1 using the transformation (expression + 2) × 8, and set to zero the expression of SPI1, IKZF1, IKZF3, TCF3, EBF1, FLT3, and IL7R. To drive differentiation toward CLP and related lymphoid lineages, we apply the opposite pattern: up-regulating SPI1, IKZF1, IKZF3, TCF3, EBF1, FLT3, IL7R, DNTT, LYL1, and LMO2, while down-regulating GATA1, GATA2, CEBPA, CEBPB, GFI1, and CSF1R. These genes are selected based on well-established roles in hematopoietic fate specification reported in previous studies. Each perturbed cell is then used as an input to CellTempo to generate corresponding developmental trajectories.

To evaluate differentiation outcomes, we classify all generated cells using CellTypist models trained on the original dataset. To map model-generated cells back onto the potential landscape, we embed both real and generated cells into the pretrained VQ-VAE latent space, construct a PCA representation using the latent embeddings of the real cells, and project the generated cells into this same PCA space. For each generated cell, we identify its nearest neighbor among the real cells in PCA space and use the corresponding real cell as its representative location on the potential landscape. Plotting these mapped points enable us to compare natural and perturbed developmental trajectories directly on the landscape and to observe how gene perturbations alter the paths that cells follow across the fixed potential surface.

### Fine-tuning CellTempo for chemical perturbation prediction

Predicting cellular responses to chemical perturbations is another key challenge in single-cell research, with important implications for drug discovery. To enable chemical perturbation– response prediction, we fine-tune the pretrained CellTempo model on paired control– perturbation data. During fine-tuning, the model is conditioned on a control cell together with drug identity and dosage information, and learns to generate the corresponding perturbed cell state.

We first construct the control–perturbation pairs. For each perturbation condition in the dataset, we select control cells originating from the same cell line and same plate to ensure matched experimental background. We then randomly sample one control cell and one perturbation cell to form a training pair (**x**_ctrl_, **x**_pert_). Both cells are encoded using the pretrained VQ-VAE encoder to obtain discrete code sequences:

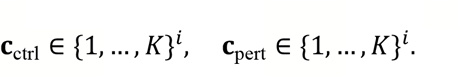

We then formulate the input token sequence for finetuning. For each pair, we construct a perturbation sentence that embeds both biological and experimental metadata. The final tokenized sequence is:

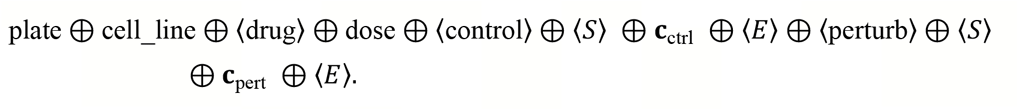

Where plate is the plate ID, cell_line is the cell line name, ⟨drug⟩ is a placeholder for drug SMILES, dose is the drug dosage, ⟨control⟩ and ⟨perturb⟩ are the control cell and perturbed cell indicators. This sequence is concatenated into a single trajectory-like sentence and used as input to the Transformer during fine-tuning.

To generalize to unseen drug molecules, we need to obtain universal molecular representations. During the forward pass, we replace the embedding of the drug token ⟨drug⟩ using a molecular representation computed by UniMol2^46^. Given a SMILES string *s*, UniMol2 produces a molecular embedding with dimension *d*_*m*_:

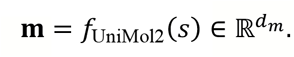

This embedding is projected into the model’s token embedding dimension *d* via a linear layer:

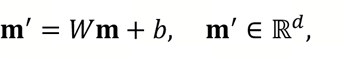

and used to replace the learned embedding of ⟨drug⟩. Dose information is encoded as a bucketized categorical token and embedded into the same vector space. This design allows the Transformer to jointly attend to the control cell’s discrete representation, the chemical structure of the drug, cell-line identity and plate metadata, and dosage information.

We fine-tune the pretrained CellTempo model using an autoregressive objective identical to the pretraining procedure. The model receives the entire perturbation sequence and is trained to predict the perturbed cell’s tokens **c**_pert_ using causal masking. Let the full token sequence be (*w*_1_, …, *w*_*L*_). The training loss is the cross-entropy:

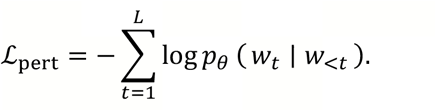

Only tokens corresponding to the perturbed cell are included in the loss to avoid over-constraining metadata tokens.

At inference time, we provide the model with the control cell code sequence **c**_ctrl_, metadata tokens (plate, cell line), drug embedding **m**′, and dose token. The model then autoregressively generates **ĉ**_pert_ representing the predicted perturbed cell in the discrete latent space. Finally, the VQ-VAE decoder reconstructs the full gene-expression profile:

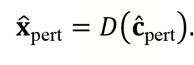

We fine-tuned the model on the Tahoe-100M perturbation dataset. To evaluate generalization, we constructed two disjoint test sets: (i) an unseen cell-line test set, in which cell lines do not appear during training but drugs are known; and (ii) an unseen perturbation test set, in which drug perturbations are novel while cell lines are known. These splits assess CellTempo’s ability to generalize across new biological contexts and new chemical perturbations, respectively.

Across both settings, CellTempo-Pert matched or outperformed established baselines, including PRNet, a linear model, and a multilayer perceptron (MLP) model (Figure S4B–C), indicating that the CellTempo architecture effectively captures immediate drug-induced state transitions. The details about baseline models and the evaluation metrics can be found in below paragraphs.

### Baseline models for chemical perturbation prediction

#### Linear

The Linear baseline models perturbation effects using a simple linear regression framework. Given a control cell with gene-expression vector 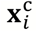 and Functional-Class Fingerprints (FCFP) of the drug SMILES 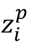 generated by RDKit^60^, the model predicts the perturbed expression profile through a linear transformation:

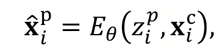

where *E*_*θ*_ is a fully connected layer parameterized by weight matrix *θ*. The model is trained to minimize mean squared error between the predicted and observed perturbed expression:

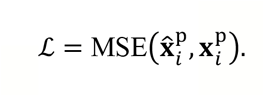

#### MLP

The MLP baseline extends the linear model by incorporating nonlinear transformations through a small feed-forward neural network. The input, comprising both the control-cell expression 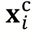 and perturbation drug’s FCFP 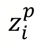, is passed through a multilayer perceptron:

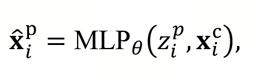

where *θ* denotes the network parameters. The network architecture consists of an input layer, a linear layer with output dimension 128, BatchNorm1d, a LeakyReLU activation, a final linear layer projecting back to gene-expression dimensionality, and a ReLU activation. The model is optimized using the same MSE objectiveas the linear baseline.

#### PRNet

PRNet is a perturbation-conditioned deep generative model designed to predict transcriptional responses to chemical perturbations, including novel compounds not seen during training^50^. PRNet explicitly models how a cell’s gene-expression profile changes under a perturbation by jointly conditioning on the unperturbed transcriptional state and compound structure. In this model, each input consists of an unperturbed gene-expression vector and the FCFP representation of the perturbing compound derived from the RDKit. PRNet embeds chemical structure information and the dosage information into a learned latent space via a Perturb-adapter, enabling generalization to unseen compounds. A Perturb-encoder then integrates this chemical embedding with the baseline expression profile to capture the compound’s effect on cellular transcriptional programs, and a Perturb-decoder generates the corresponding perturbed expression profile.

### Metrics for evaluating chemical perturbations

To quantitatively evaluate the accuracy of predicted chemical perturbation responses, we assess model performance at both the gene level and the population (pseudobulk) level using a set of complementary metrics commonly adopted in perturbation-response prediction.

#### Differentially expressed gene (DEG) direction agreement

This metric evaluates whether the model correctly predicts the direction of differential expression for genes significantly perturbed in the ground truth. Let 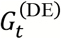 denote the intersection of ground-truth and predicted DEGs for a perturbation *t*. For each gene 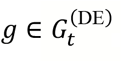, let Δ_*tg*_ and 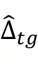 denote the true and predicted log-fold changes. Directionality agreement is defined as the fraction of genes whose change directions match:

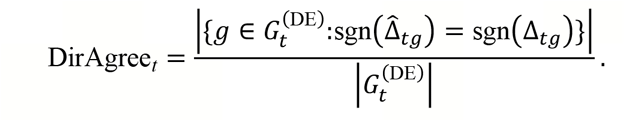

This metric measures whether the model predicts up- and down-regulation trends correctly.

#### DEG Fold-change SCC

For the same DEG set 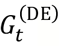, this metric computes the SCC between predicted and true log-fold changes. It evaluates whether the overall ordering of gene responses is preserved.

#### Pseudobulk delta PCC

At the pseudobulk level, we compute the mean expression profile of perturbed cells 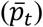 and control cells 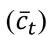, and define the expression delta:

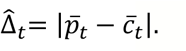

The PCC between predicted detla 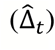 and observed delta (Δ_*t*_) assesses whether the model captures the magnitude of perturbation effects.

#### Pseudobulk Reconstruction Error (MAE)

To measure absolute deviation between predicted and observed pseudobulk profiles, we compute:

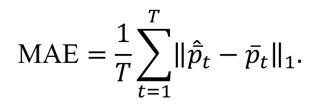

These metrics quantify reconstruction fidelity of global perturbation effects.

#### Discrimination score L2

This metric evaluates whether the predicted response for perturbation *t* is closest to the correct ground-truth profile among all perturbations. Let *d*(⋅,⋅) be the L2 distance metric. The ranking score:

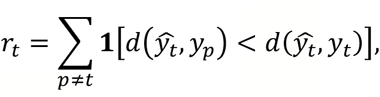

counts how many incorrect perturbations appear more similar to the prediction than the true target. Here **1** is an indicator function, true returns 1 and false returns 0. We report the normalized inverse score:

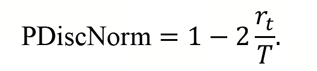

A random predictor scores ≈ 0, while an ideal predictor scores 1.

#### Energy-distance PCC

We quantify how well the model recovers the magnitude of perturbation-induced shifts in the cell-state distribution using an energy-distance–based score. For each perturbation *t*, given samples *X*_*t*_ = {*x*_*i*_} from the perturbed population and *Y* = {*y*_*j*_} from the corresponding control population, we approximate the energy distance between the two distributions as

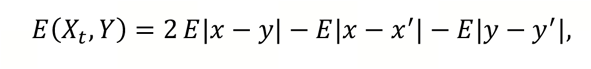

where expectations are estimated by the mean pairwise Euclidean distances between all perturbed–control pairs, perturbed–perturbed pairs, and control–control pairs, respectively. For each perturbation *t*, we compute this quantity both for the ground-truth data, 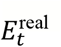, and for the model predictions, 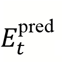, reusing the control–control term across perturbations. The final evaluation score is the PCC across perturbations:

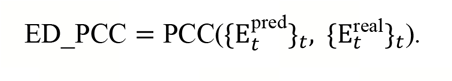

A high ED_PCC indicates that the model accurately reproduces the relative strengths of perturbation effects across different perturbations, even if the exact gene-level profiles are not perfectly matched.

### Forecasting OPC-directed differentiation from iPSC-derived glial progenitors

#### Dataset and starting cell population

We evaluated CellTempo using a published longitudinal single-cell RNA-sequencing dataset tracking the differentiation of human iPSCs into mixed populations of neurons, astrocytes, and oligodendrocyte-lineage cells ^45^. The day-30 proliferative glial progenitor cells (GPCs) were selected as the starting population for all OPC-directed forecasting experiments. The same set of starting cells was used for all control and perturbation conditions.

#### Selection and *in silico* activation of OPC-associated perturbation genes

To identify candidate genes associated with OPC differentiation, we integrated lineage association and differential expression analyses following the pipeline of CellRank^63^. A diffusion map was first constructed using Diffusion pseudotime^64^. For each cell, an OPC fate score was calculated as the cosine similarity between its position and the centroid of the OPC_main population in the diffusion-map space. For each gene expressed in more than 5% of cells, we calculated the Pearson correlation between its log1p-normalized expression and the OPC fate score, denoted as PCC_OPC_. Differential expression analysis between OPC_main and astrocyte cells was performed using the Wilcoxon rank-sum test implemented in Scanpy. Candidate genes were ranked using a composite score integrating their association with OPC fate, differential expression magnitude (logarithmic fold change), and statistical significance (adjusted *p*-value):

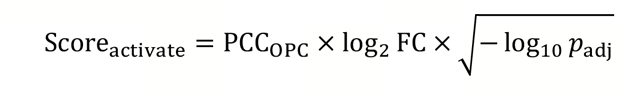

Genes satisfying PCC_OPC_ > 0.3, log_2_ FC > 1, and adjusted *p*-value < 0.05 were considered candidate activators of OPC differentiation. We selected prioritized all transcription factors using Score_activate_, resulting in the final gene set, including OLIG1, PTN, ETV1, DLL3, C2orf27A, LIMA1, SCRG1, SMOC1, PTPRZ1, SIRT2, OLIG2, and PDGFRA. The perturbed cells were subsequently used as the initial states for trajectory forecasting by CellTempo.

The selected genes were activated simultaneously in the day-30 proliferative GPCs. For each selected gene, its expression value was transformed according to x_new_ = 2x_orig_ + 8, where x_orig_ and x_new_ represent expression values before and after perturbation, respectively. The same transformation was applied across all starting cells and selected genes. The perturbed and unperturbed cells were subsequently used as initial states for trajectory forecasting by CellTempo.

#### Compound selection and construction of multi-compound perturbation signatures

For single-compound analyses, we selected compounds used in a published patent^48^ as candidate OPC-promoting compounds, together with common compounds expected to have little or no effect on OPC differentiation as controls. The candidate compounds included TTNPB, SC79, MHY1485, purmorphamine, XAV939, LDN193189, Go6983, MK2206, AZD3147, and AICAR, whereas the control compounds included aspirin, caffeine, glycerol, urea, acetamide, propylene glycol, ibuprofen, benzene, ethanol, and ethylene glycol. DMSO was used as the vehicle reference. We further evaluated the multi-compound iPSC-to-OPC induction strategy described in the published patent. The protocol consisted of TTNPB, SC79, MHY1485, purmorphamine, XAV939, LDN193189, Go6983, and MK2206.

To evaluate candidate chemical interventions, we first used CellTempo-Pert to predict the immediate transcriptional responses of day-30 proliferative GPCs. For each compound, CellTempo-Pert independently generated the predicted cell states following treatment with the compound and with DMSO.

Compound-specific transcriptional responses were identified by comparing the compound-predicted state with the DMSO-predicted state. Differential expression analysis was performed using the Wilcoxon rank-sum test, and genes were selected according to log2FC values. The selected upregulated and downregulated genes were then applied as signed *in silico* perturbations to the original day-30 proliferative GPCs. Upregulated genes were modified according to x_new_ = 2x_orig_ + 8, whereas downregulated genes were set to zero. The resulting perturbed cells were used as starting states for long-term trajectory forecasting by CellTempo.

For the multi-compound strategy, the immediate transcriptional response to each constituent compound was inferred independently using CellTempo-Pert relative to its matched DMSO-predicted state. The resulting compound-specific signatures were combined as a union to construct an aggregate perturbation signature representing the predicted effects of the protocol components. Each gene in this union is perturbed once, with upregulated genes modified according to x_new_ = 2x_orig_ + 8 and downregulated genes set to zero.

#### Cell-type annotation of forecast trajectories

Cell identities along generated trajectories were assigned using marker-based scoring. The score_genes function implemented in Scanpy was used to calculate a score for each cell and each candidate cell type based on curated marker-gene sets. Cells were assigned to the cell type with the highest marker score.

Marker genes were selected from the original study^45^ and defined as follows: neuron, MAP2, TUBB3, SYP, DCX, RBFOX3, and SNAP25; NPC/GPC, SOX2, NES, VIM, PAX6, HES1, and FABP7; proliferative GPC, MKI67, TOP2A, PCNA, TUBA1B, HMGB1, HMGB2, H2AFZ, TYMS, PCLAF, and MAD2L1; astrocyte precursor, GFAP, S100B, ALDH1L1, AQP4, SLC1A3, and CD44; astrocyte, GFAP, S100B, ALDH1L1, AQP4, GJA1, and APOE; OPC, PDGFRA, OLIG2, SOX10, CSPG4, OLIG1, and NKX2-2; pre-OPC, PDGFRA, OLIG1, GPR17, BCAS1, and NKX2-2; and oligodendrocyte, MBP, MOG, PLP1, MAG, CNP, and MOBP.

OPC-lineage abundance at each prediction step was calculated as the proportion of generated cells annotated as OPC and pre-OPC.

## Data availability

scBaseTraj is available on https://huggingface.co/datasets/EperLuo/scBaseTraj. The human bone-marrow hematopoiesis dataset^30^, mouse pancreatic endocrine differentiation dataset^23^, and erythroid differentiation during mouse gastrulation^24^ dataset are obtained from the scVelo Python package^17^. The human uterus dataset^25^ (SRX12173016 and SRX12173003) and immune cell dataset^65^ (SRX11980210) are selected from the scBaseCount database^8^.The OPC induction data^45^ are available in the Gene Expression Omnibus (GEO) repository under accession GSE211140.

## Code availability

CellTempo code, documentation, and tutorials are available at https://github.com/EperLuo/CellTempo.

### Acknowledgements

The work is sponsored in part by the National Key R&D Program of China (2025YFC3409300, 2024YFF0729200), and National Natural Science Foundation of China (92470105, 62373210).

## Supplementary information

**Figure S 1.**
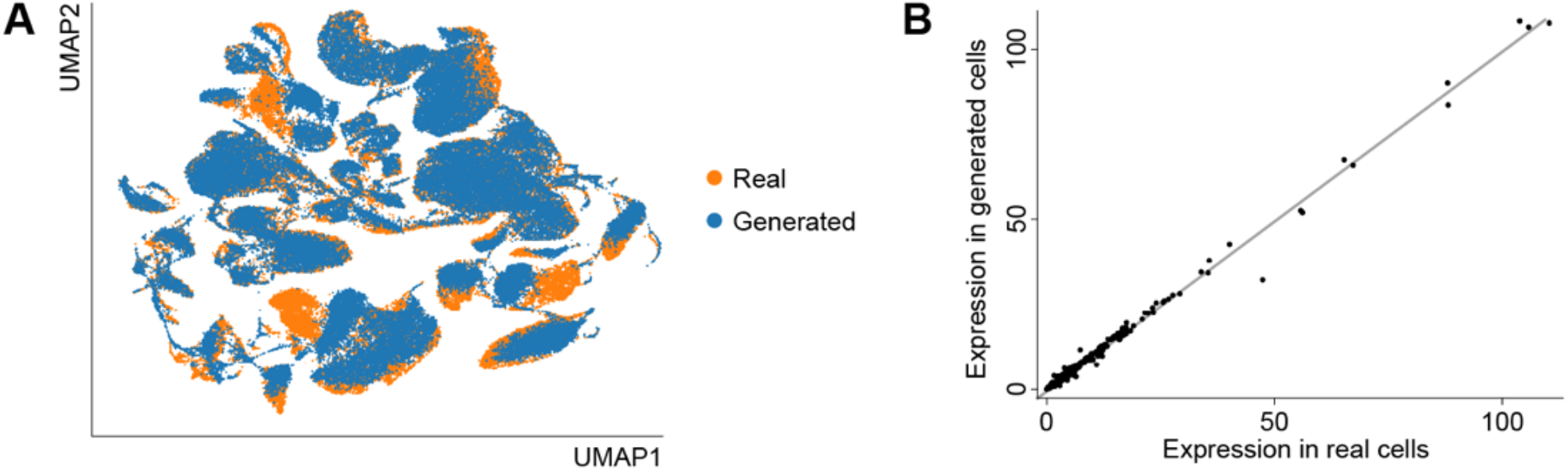
Reconstruction quality of the cell VQ-VAE. (A) UMAP of real cells in the test dataset and cells reconstructed by the cell VQ-VAE. (B) Average gene expression of real cells in the test dataset and cells reconstructed by the cell VQ-VAE. Each point represent a gene. The gray line denotes perfect agreement between real and generated expression.

**Figure S 2.**
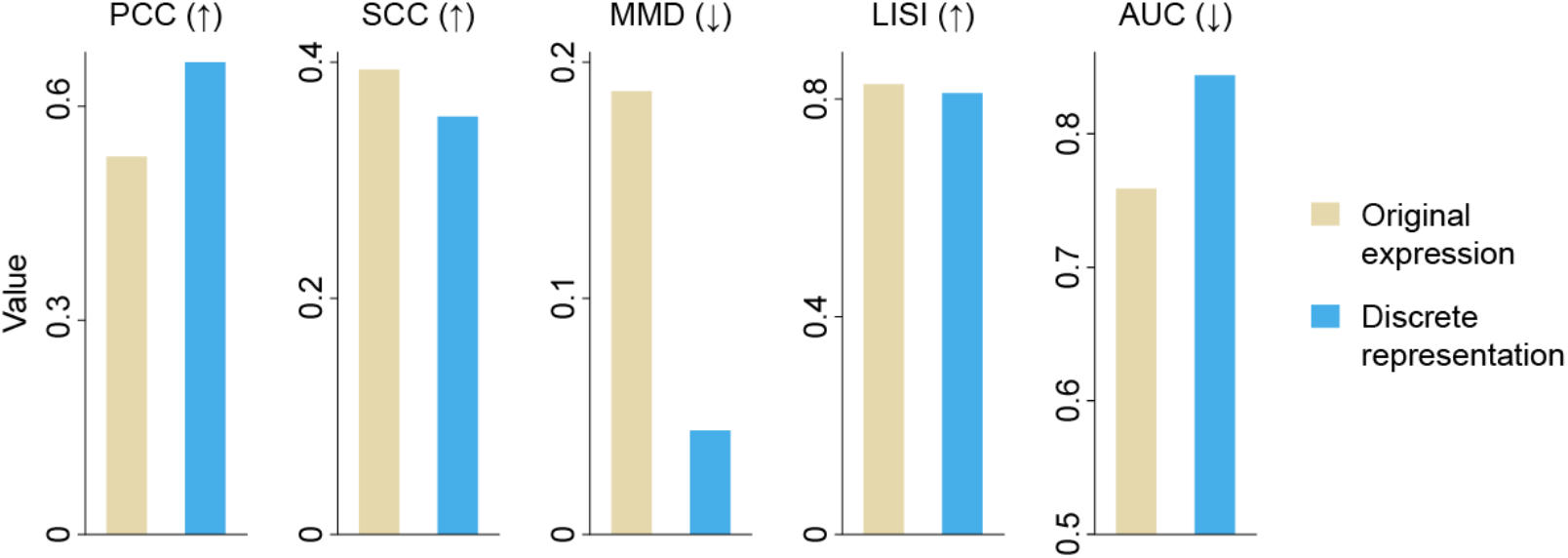
Performance comparison between latent-space and full-expression-space models. Upward arrows indicate that higher values correspond to better performance, whereas downward arrows indicate the opposite.

**Figure S 3.**
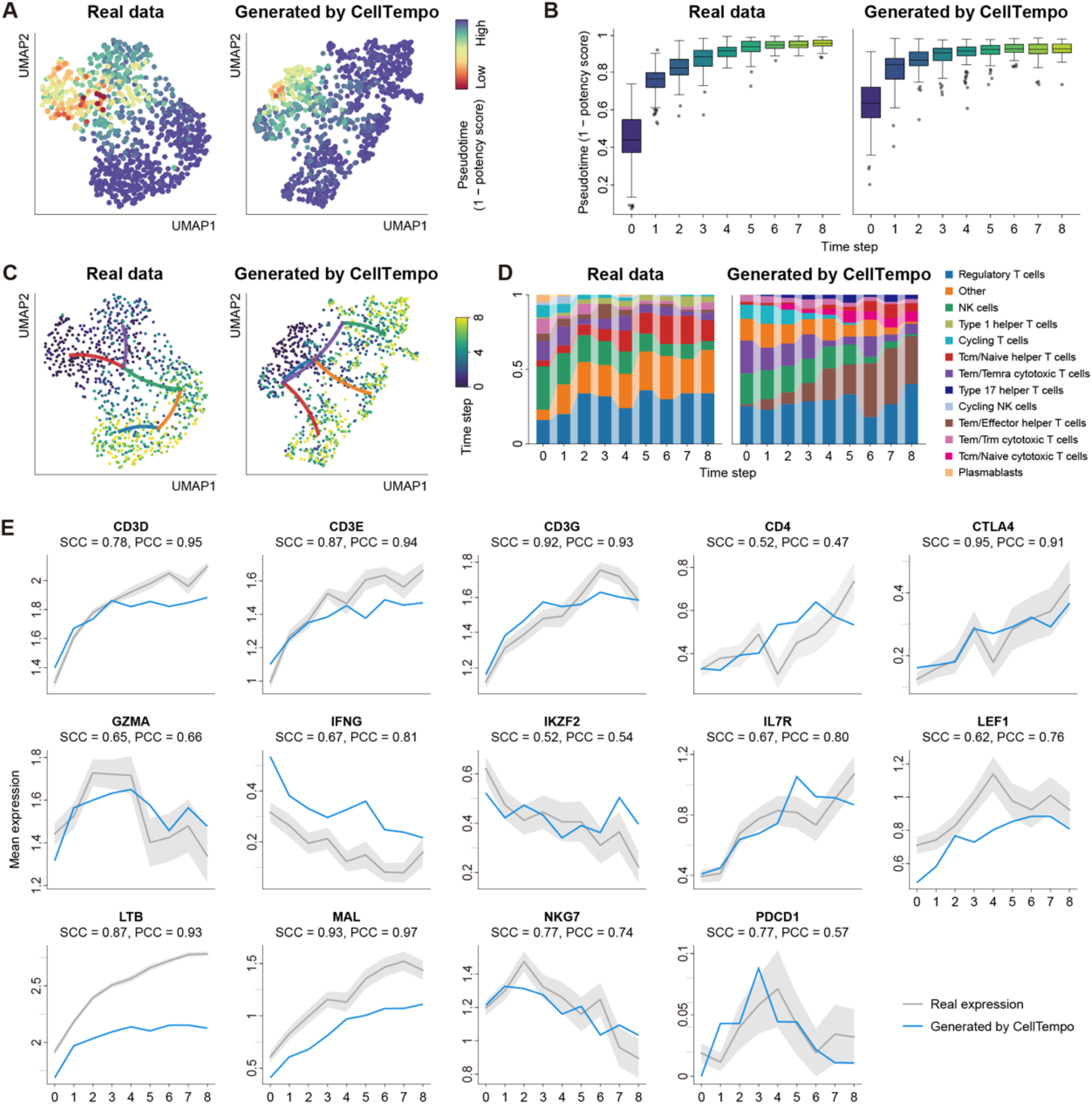
Experiment results on the human immune cell dataset. (A) UMAP of real and CellTempo-generated cells, colored by pseudotime defined as 1 – CytoTRACE2 potency score. (B) Pseudotime of real and generated cells at each time step. (C) UMAP of real and generated trajectories colored by time step. Lines represent high-probability transitions between clusters based on a PAGA-derived cluster transition graph. (D) Cell-type composition at each time step in real and generated trajectories. (E) Temporal dynamics of marker gene expression across time steps

**Figure S 4.**
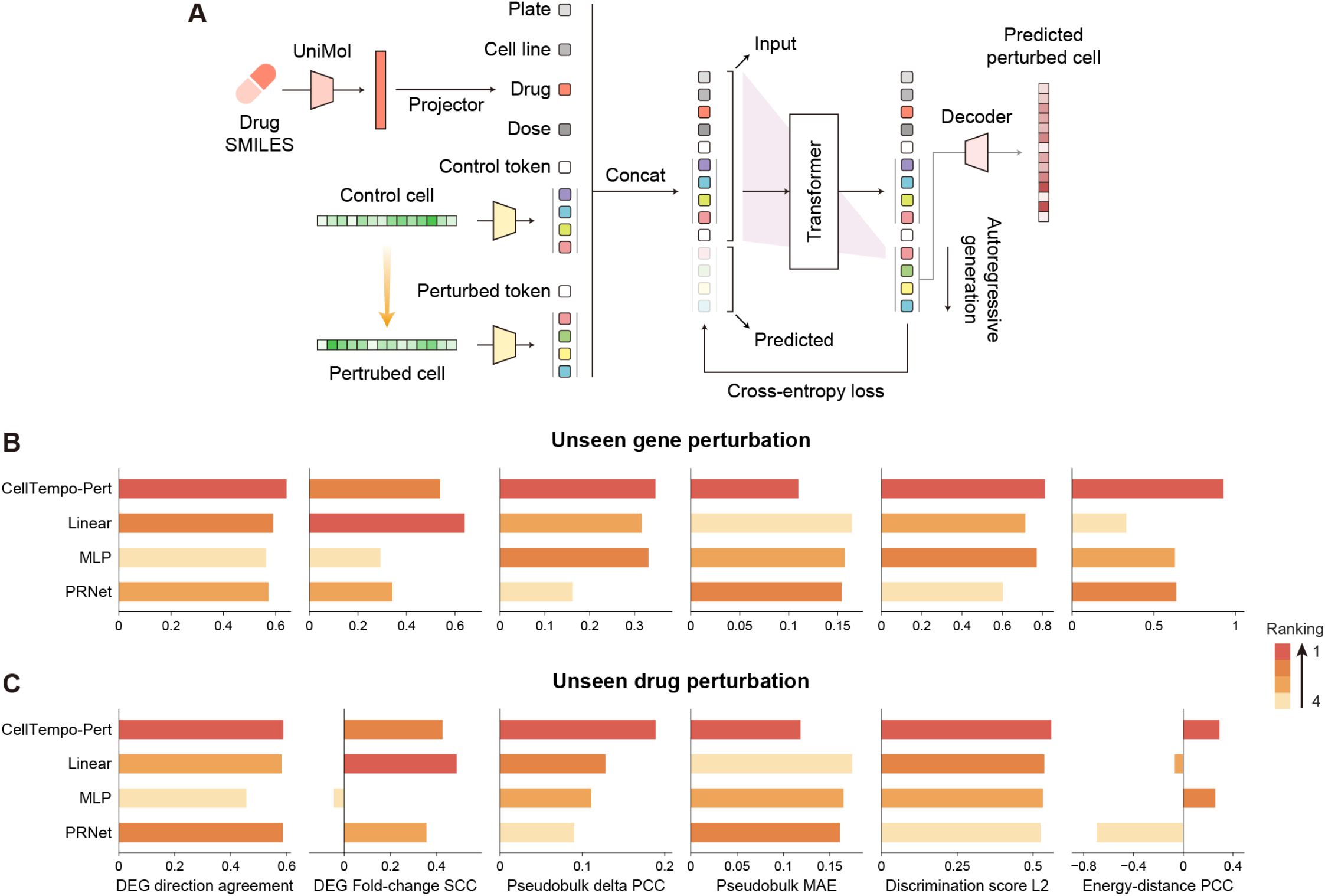
CellTempo-Pert for drug perturbation prediction. (A) We model each control–perturb cell pair as a two-step trajectory, combined with plate, cell line, drug, and dose information for fine-tuning CellTempo to derive CellTempo-Pert. Drug representations are obtained from a pretrained UniMol2 model. Conditioned on this meta-information and control-cell tokens, CellTempo autoregressively predicts the perturbed-cell tokens. The final perturbed-cell gene expression profile is reconstructed using the cell VQ-VAE decoder. (B-C) Performance of CellTempo-Pert and other methods in unseen cell-line perturbations (B) and unseen drug perturbations (C).

